# Development of an image-guided non-vitrectomy subretinal access approach for trans-scleral cell and gene therapy delivery

**DOI:** 10.1101/2025.02.25.640217

**Authors:** Mandeep S. Singh, Shoujing Guo, Christopher B. Toomey, Minda McNally, Sarah Harris-Bookman, Kang Li, Dzhalal Agakishiev, Shuwen Wei, Soohyun Lee, Kathleen Perrino, Kevin C. Wolfe, Jin U. Kang

**Author notes:** Joint last authors (Kevin C. Wolfe, Jin U. Kang). **Corresponding author:** Mandeep S. Singh, MD, PhD Wilmer Eye Institute, Johns Hopkins University 600 N. Wolfe St Baltimore, MD 21287.

## Abstract

**Purpose:** Regenerative therapies for retinal diseases include cell and gene therapy modalities that are targeted to the subretinal space. Several recent clinical trials have shown that the morbidity of surgical access is the major limitation of safe subretinal space delivery. We aimed to develop an image-guided procedure for minimally invasive subretinal access (MISA) as a platform to deliver therapeutic agents for the treatment of degenerative retinal diseases.

**Methods:** We engineered prototypes of a novel common-path swept source optical coherence tomography (CP-SSOCT)-enabled needle, coaxial guide (COG), and subretinal access cannula (SAC). We pilot tested the MISA procedure in *ex vivo* bovine eyes and *in vivo* porcine ocular surgery.

**Results:** A- and M-mode scan recordings of *ex vivo* and *in vivo* animal eye models demonstrated that CP-SSOCT imaging from the scleral side (*ab externo*) was capable of identifying the retinal laminae and the sub-retinal space. We show results from *in vivo* porcine MISA surgeries (N=4) using the novel CP-SSOCT-enabled sub-retinal injection needle, COG, and SAC through the transscleral approach. The MISA approach enabled subretinal device placement in the posterior pole, however, cases of retinal incarceration and retinal perforation were encountered.

**Conclusions:** We describe a novel CP-SSOCT-guided subretinal access approach that, with further optimization, may be useful in regenerative retinal surgery.

## Introduction

Acquired or inherited degenerative retinal diseases, including non-exudative age-related macular degeneration (AMD), retinitis pigmentosa (RP), and Stargardt macular dystrophy, are major global causes of blindness. The cells that degenerate in these conditions are the photoreceptor and retinal pigment epithelium (RPE) cells. Electronic, stem cell, and gene therapy modalities are in development, aiming to preserve, restore, or replace the structure and/or function of the photoreceptor and RPE cell layers[1, 2]. These modalities are delivered to the anatomical space known as the subretinal space.

The subretinal space is a potential space that is located between the photoreceptor and RPE cell layers that are normally adherent to one another[3]. The actual subretinal space is created by the pathologic or iatrogenic accumulation of material between the photoreceptor and RPE cell layers. In various disease states, exudate or hemorrhage can accumulate in the subretinal space[4]. In vitreoretinal surgery, the subretinal space can be created by infusing saline between the photoreceptor and RPE cell layers to create a space for the delivery of therapeutic agents. Soluble subretinal medications that are targeted to the subretinal space include tissue plasminogen activator and voretigene neparvovec-ryzl. The former is used to liquefy submacular hemorrhage[5] and the latter for gene therapy of *RPE65*-associated retinopathy[6]. The alpha-IMS electronic retinal implant, a microphotodiode array to replace the function of lost photoreceptor cells, was also designed for subretinal placement. Cell therapy agents, including human umbilical tissue-derived cells (hUTC)[7] and stem cell-derived RPE cells[8–12] are also being targeted to the subretinal space.

Conventionally, the transvitreal *ab interno* approach has been used to gain access to the subretinal space. The key steps of this approach are removing the vitreous gel (i.e., vitrectomy) and inducing a perforation in the retina (i.e., retinotomy). Complications of *ab interno* access have included cataract formation, retinal detachment, intraocular hemorrhage, proliferative vitreoretinopathy (PVR), and epiretinal membrane (ERM) formation[8, 9, 13–16]. Another important risk of *ab interno* access is the potential for the retrograde reflux of therapeutic agents into the vitreous cavity via the retinotomy[17], thus reducing the subretinal therapeutic dose and promoting antigen exposure that may heighten the risk of immune rejection[11]. Taken together, delivery-related complications of the *ab interno* approach have the potential to reduce efficacy, trigger adverse events, and threaten potential visual gains, thus compromising treatment outcomes. The vitrectomy and retinotomy steps are the main sources of the surgical morbidity of the *ab interno* approach.

The observations above frame the unmet need to develop a subretinal delivery solution that minimizes surgical morbidity by avoiding vitrectomy and retinotomy. To address this need, we developed a novel approach termed minimally invasive subretinal access (MISA). MISA uses an image-guided *ab externo* route to the subretinal space. Reflecting widespread recognition of the potential advantages this approach, several *ab externo* devices for delivery to the suprachoroidal space[18–22] and subretinal space[7, 23, 24] have been developed, although these have lacked image guidance thus far.

To increase surgical precision and safety of the MISA approach, we leveraged the technical advance of endoscopic common-path swept-source optical coherence tomography (CP-SSOCT) imaging. CP-SSOCT imaging enables directional, real-time, and depth-resolved A-scan visualization of structures distal to and coaxial with the distal sensor[25–27]. We incorporated a CP-SSOCT distal sensor to develop MISA according to the principles of image-guided surgery (IGS): to use imaging technology in the operative field to provide real-time information on the precise location of a surgical instrument relative to anatomic structures of interest.

The observations reported here reflect the relatively modest goals of the pilot experiments that we conducted using prototypes of the components described below. Because of the lack of prior data regarding this approach, we found it necessary to perform these pilot studies to validate conceptual feasibility, and identify technical risks, prior to launching a full-scale study to obtain surgeon feedback, iterate the prototype designs, and test optimized system performance in a larger cohort of animals. Here we present data from those pilot experiments that demonstrate proof-of-concept of the MISA approach as a novel strategy in regenerative retinal surgery.

## Materials and Methods

### MISA system components

The components of the MISA system were the CP-SSOCT, the coaxial guide (COG), and the subretinal access cannula (SAC).

#### CP-SSOCT

A CP-SSOCT device was designed to acquire real-time depth information of the imaged tissue and the delivery device position. The system utilizes a reference signal from the distal end of the fiber probe. The sample and reference beam share the same single-mode fiber, thus limiting dispersion and polarization noise. The light source was a swept-source OCT engine (AXSUN, Billerica MA USA) with a center wavelength of 1060 nm and a sweeping rate at 100 kHz. A broadband circulator (OF-Link, BPICIR-1060-H6) was used to combine the reference and sample beam to generate the interference signal. The spectral data was then detected by a balanced detector integrated into the OCT engine and collected by a frame grabber (National Instrument, PCI-E-1433). The overall system was packed in a benchtop electronics enclosure (**Figure 1A**). The measured axial resolution was 4.5 microns inside the tissue and the axial imaging range was 3.7 mm in the air. A CP-SSOCT distal-sensor guided injection device was created by integrating a single-mode optical fiber as the optical sensor with a high-index epoxy lens, secured inside a 30-gauge needle with a fixed offset from the tip[27]. A three-way stopcock facilitated an additional port for a syringe to be connected for injection of material. The needle and stopcock were mounted on a translation stage (**Figure 1B**). An articulated arm was used to enhance mobility.

**Figure 1.**
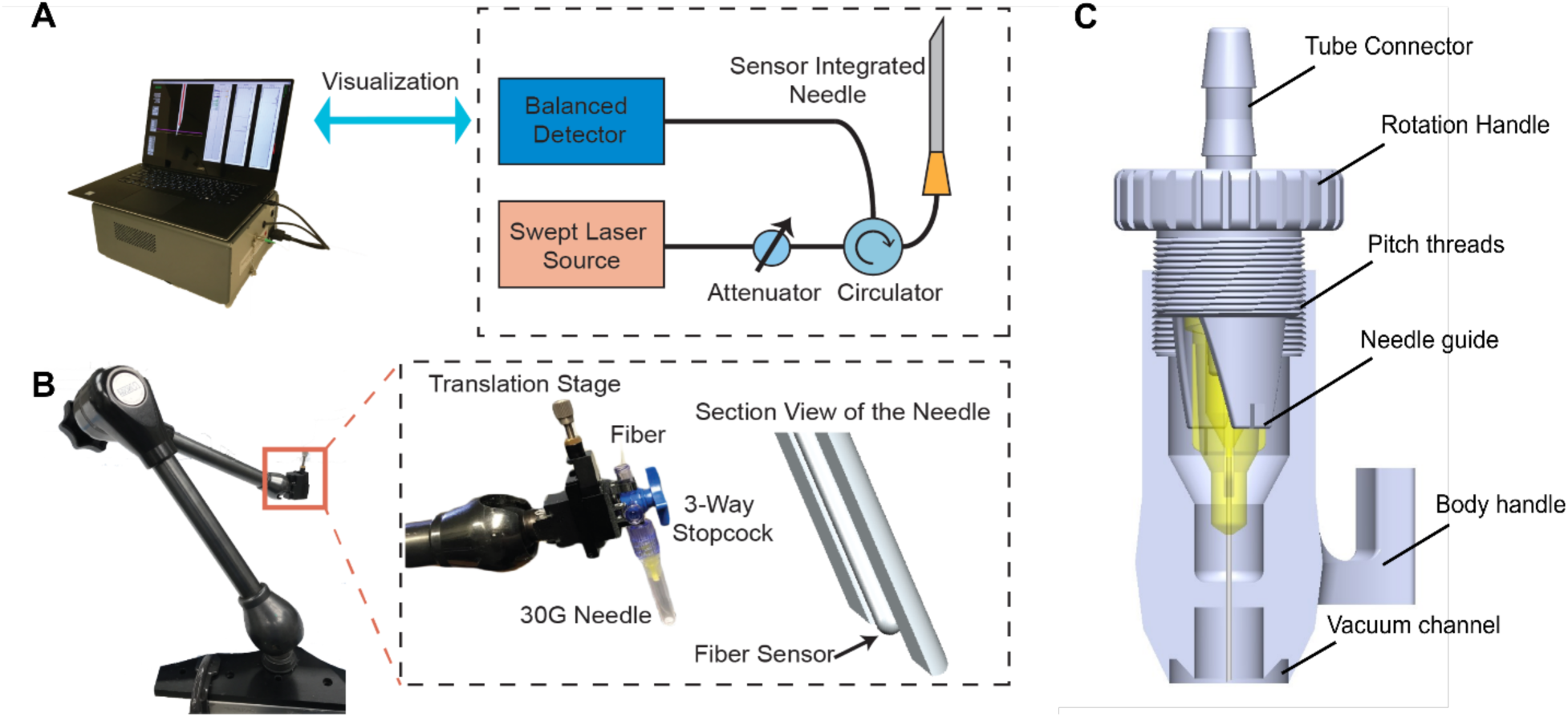
Schematic of the common-path swept source optical coherence tomography (CP-SSOCT) and coaxial guide (COG) components of the minimally invasive subretinal access (MISA) system. (A) CP-SSOCT device diagram showing the balanced detector, swept source laser source, attenuator, and circulator connected to the sensor integrated needle. (B) CP-SSOCT distal-sensor guided injection device with articulated arm supporting and stabilizing the translation stage, and (C) COG device that enabled secure coupling the sensor integrated needle and driver to the scleral surface by vacuum suction.

### Coaxial guide (COG)

The COG was made to stabilize the CP-SSOCT device intraoperatively by vacuum suction on the scleral surface (**Figure 1C**). The COG consisted of two parts: the eye adaptor and the needle drive. The eye adaptor formed the main body of the COG and it included a vacuum channel (diameter of 10 mm) to fix the device onto the eye. The needle drive had a thread pitch of 500 microns for advancing the needle. Rotation of the handle advanced the needle forward. The COG was 3D-printed by a Stratasys Objet30 Pro PolyJet printer using VeroBlue material.

### Subretinal access cannula (SAC)

The intent of the SAC was to create a smooth-gliding passageway for therapeutic material to be inserted into the subretinal space. For the purposes of this project, the envisioned therapeutic material was a construct of the approximate size, shape, and stiffness of a preformed sheet of cells-with-scaffold similar to those used in several recent clinical trials[9]. Therefore, the SAC lumen had to be at least 5.00 mm wide. The SAC was made of top and bottom polyimide layers encased in a latex tube (**Figure 2**). The latex tube was Mehron liquid latex mixed with distilled water (3:1 ratio). The latex mixture was degassed and the tubes were created by dip coating 3 mm diameter glass tubes which were cured at 50 °C for 12 hours. The tubes were removed and rinsed with soap. The composite tube was created by encasing top and bottom polyimide layers in the latex tube. The latex tube started at the shoulder of the tip and extended to 10 mm from the proximal end. A flexible cyanoacrylate adhesive was used to adhere the distal ends of the polyimide layers to the latex and a primer was used to improve bond quality.

**Figure 2.**
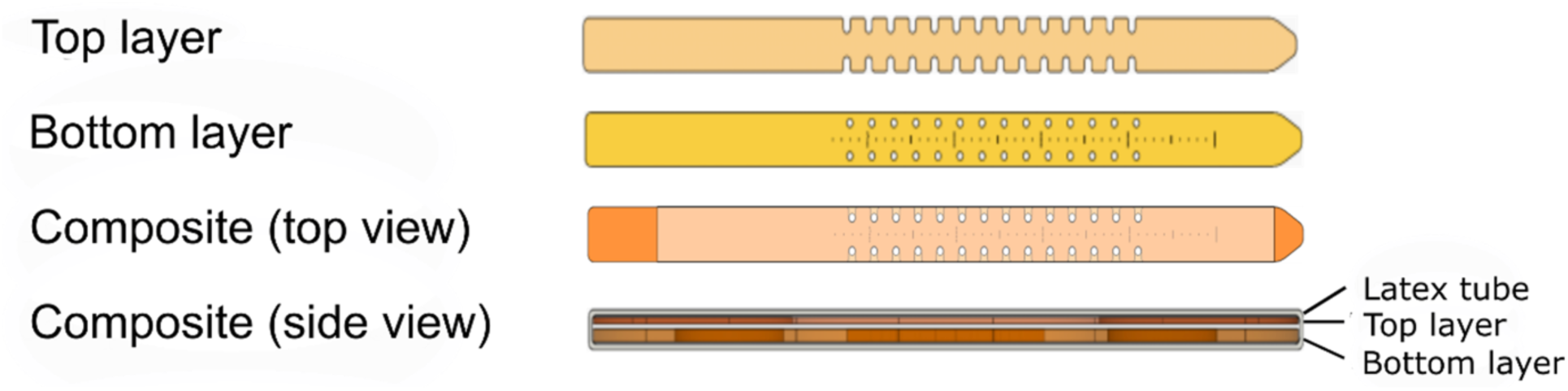
Subretinal access cannula (SAC) design. The SAC comprises two polyimide layers (top and bottom layers) encased in a latex tube. The indentations in the top layer and the holes in the bottom layer, in combination with the ruler measure marks, enable accurate insertion length determination and suturing of the SAC to the sclera for stabilization. segment/outer segment.

### *Ex vivo* testing

Fresh bovine eyes were fixed to a globe mount and the CP-SSOCT integrated subretinal 30g injection needle was positioned perpendicularly to the scleral surface at the ocular equator. A-scans along with M-mode recordings were acquired as the needle was advanced from outside the scleral surface into the sub-retinal space.

### Animals

Yorkshire pigs (Archer Farms Inc.) weighing 50 pounds were placed under ketamine (20-30 mg/kg)/xylazine (2-3 mg/kg) pre-anesthetic and isoflurane for maintenance with intravenous 5mL/kg/hr lactated ringer solution administered and monitored by a staff veterinarian. Drops of proparacaine 0.5%, tropicamide 1%, and phenylephrine 2.5% were instilled to dilate the pupils. Intraoperative data were captured photographically and the animals were sacrificed postoperatively by intravenous euthanasia.

### MISA procedure

In anesthetized pigs, a corneal traction suture was placed at the inferotemporal corneoscleral limbus and a rectus muscle was disinserted. To enable intraocular endo-illumination for the purposes of data recording in these pilot experiments, one 27-gauge valved cannula (Accurus vitrectomy system, Alcon, USA) was placed approximately 4 mm posterior to the limbus for the chandelier and another for balanced salt saline (BSS) infusion. The eye was then torted superonasally. The conjunctiva and Tenon capsule were excised capsule to expose bare sclera. A patrial thickness sclerectomy was performed. The following steps are depicted as a schematic in **Figure 3**.

**Figure 3.**
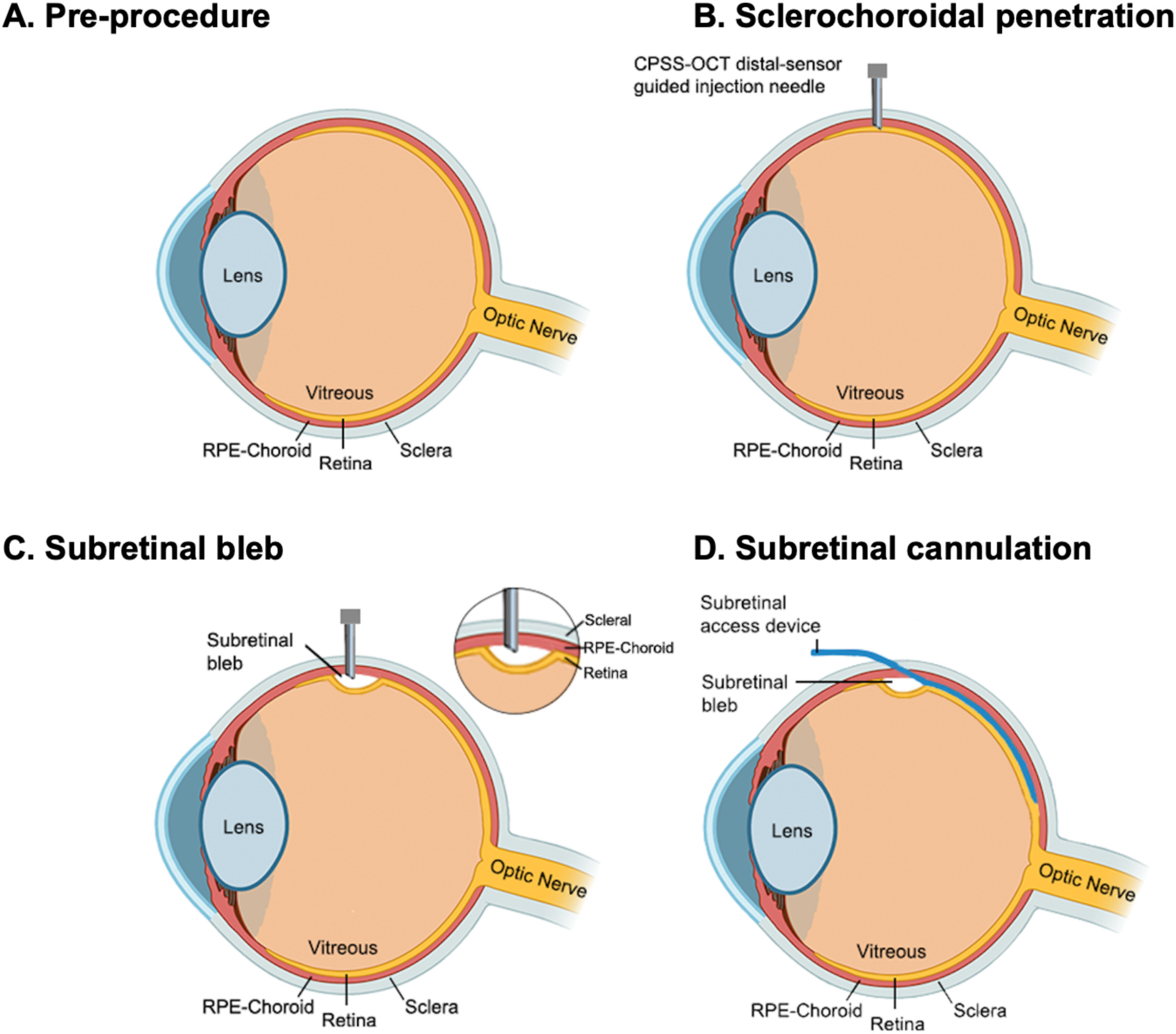
Schematic concept diagram of the minimally invasive subretinal access (MISA) procedure. This diagram shows the key steps of common path swept source optical coherence tomography (CPSS-OCT) guided transscleral subretinal access. (A) Pre-procedure ocular anatomy. (B) CPSS-OCT distal-sensor guided injection needle penetrates the sclera and choroid (and Bruch membrane, not shown) until the subretinal space is accessed. Position of the needle tip in the subretinal space is verified using CP-SSOCT. (C) A subretinal bleb is created by injection of viscoelastic material. (D) The subretinal access cannula (SAC) is inserted into the bleb and then advanced in the subretinal space to the posterior pole and macula.

#### Phase 1

A surgical pen was used to mark the position of the COG. The COG was secured over the exposed sclera by actuating the vacuum seal. The needle drive part of the COG was used to advance a 30-gauge needle (containing the OCT fiber sensor) axially towards the eye. By continued actuation of the COG needle drive, the sclera was penetrated by the needle. Real-time CP-SSOCT A- and M-mode (Quasi-B mode representation from A-scan Images recordings over time were obtained throughout the process. Once the subretinal space was identified by the *ab externo* CP-SSOCT scan, the needle was adjusted to compensate for the offset of the needle tip and the OCT fiber sensor, a subretinal bleb was created by injecting viscoelastic through the syringe port into the subretinal space. Subretinal bleb formation was verified by intraocular visualization aided by endo-illumination and scleral indentation when necessary.

#### Phase 2

To mitigate the optical effects of the thick sclera of swine, a partial thickness limbal-based scleral flap was elevated and removed with a beaver blade. A 5.5mm-wide sclerotomy was created. The exposed choroid was diathermized or laser coagulated to minimize the risk of hemorrhage and then incised with Vannas scissors to create a choroidotomy[23]. The SAC tip was inserted into the sclerotomy and choroidotomy and then advanced in the subretinal space towards the posterior pole and macula. The position of the tip of the SAC in the subretinal space of the posterior pole of the eye was verified by intraocular visualization aided by endo-illumination and intraoperative OCT. When the SAC tip was positioned in the desired location at the posterior pole, the SAC was sutured to the sclera through the premade indentations and holes to promote stability. The extraocular portion of the latex sheath was then divided using spring scissors to enable separation of the top and bottom layers of the SAC thus exposing the lumen of the SAC.

## Results

### *Ex vivo* validation of *ab externo* CP-SSOCT retinal imaging

The ability of the CP-SSOCT device to image the retinal thickness and lamination characteristics from the *ab externo* side, trans-sclerally, was tested on a bovine eye *ex vivo* (**Figure 4**). The OCT fiber sensor integrated needle was positioned perpendicularly to the scleral surface just anterior to the ocular equator as shown in **Figure 4A**. A-scan images together with M-mode images were acquired as the needle was manually advanced into the subretinal space. The locations of the choroid, RPE, photoreceptor inner and outer segment (IS/OS) line, and the inner limiting membrane (ILM) bands could be identified on the A-scan and M-mode recordings (**Figures 4B–D**), with reference to the known anatomical features of mammalian retina and the retinal lamination pattern that is typically seen with transpupillary intraocular OCT imaging. Using this system, the bovine peripheral retinal thickness measured approximately 300µm, which was comparable to that measured by a reference-based intraocular OCT imaging system (data not shown), thus further supporting our interpretation of the tissue layer correlations of the A-scan envelope. The subretinal space was identified as the space corresponding to the hyporeflective band situated between the relatively hyper-reflective RPE and IS/OS bands (**Figure 4D**).

**Figure 4.**
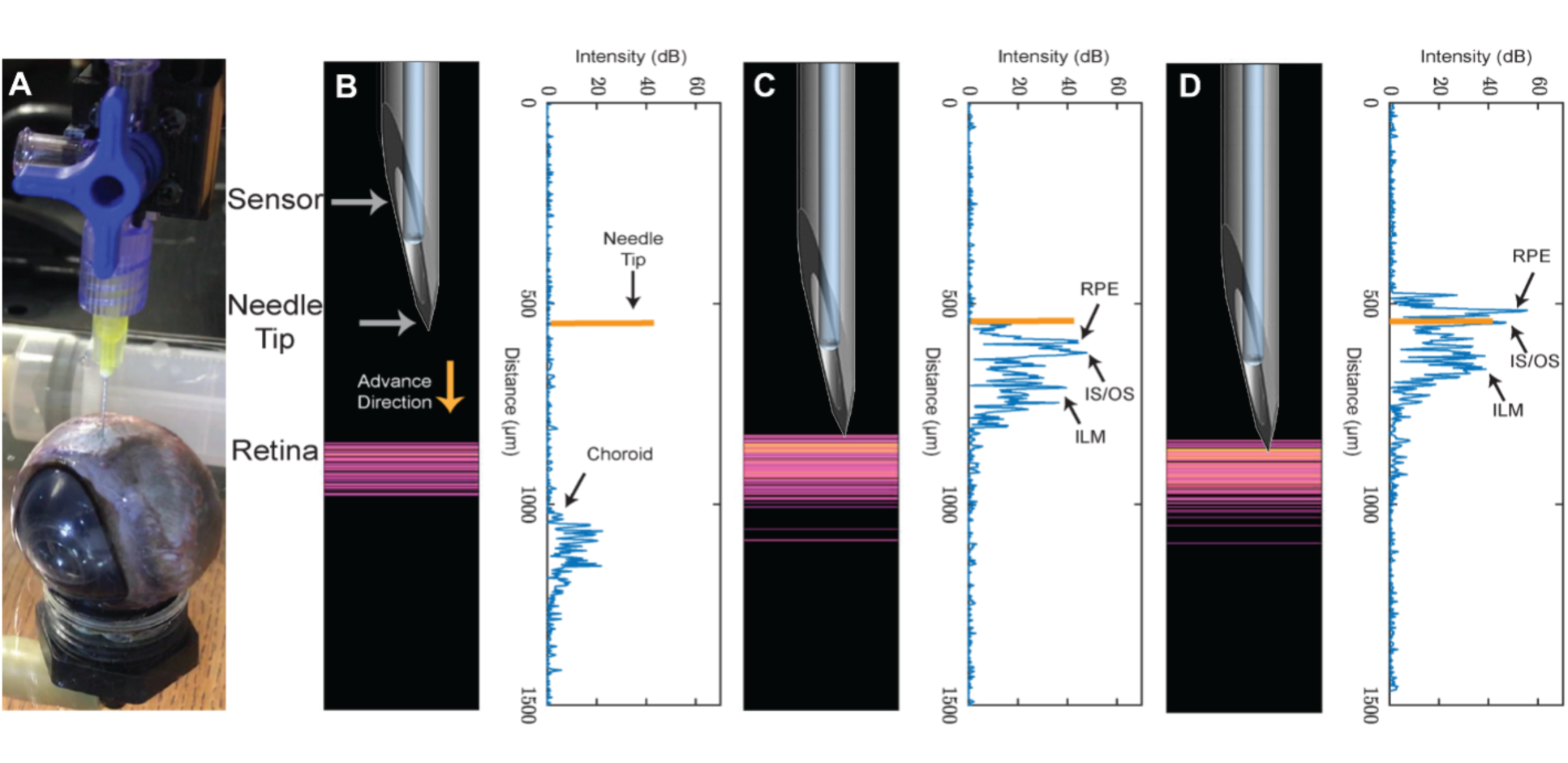
*Ex vivo* validation of common path swept source optical coherence tomography (CPSS-OCT) based detection of the subretinal plane from the *ab externo* trans-scleral direction. (A) Experimental setup for *ex vivo* testing using cadaveric bovine eyes. (B-D) A-Scan images (right) and corresponding AB-mode images (left) of bovine retinal layers as needle advances toward it, (B) ∼650 microns from choroid, (C) needle touch on the external surface of the choroid, (D) needle in the sub-retinal space. RPE: retinal pigment epithelium. IS/OS: inner segment/ outer segment. ILM: Inner retinal membrane.

### *In vivo* testing and development of MISA components

Having validated the CP-SSOCT ex vivo, we used the same system in vivo in a swine model and validated that the retinal lamination pattern was detected in this context (**Figure 5**).

**Figure 5.**
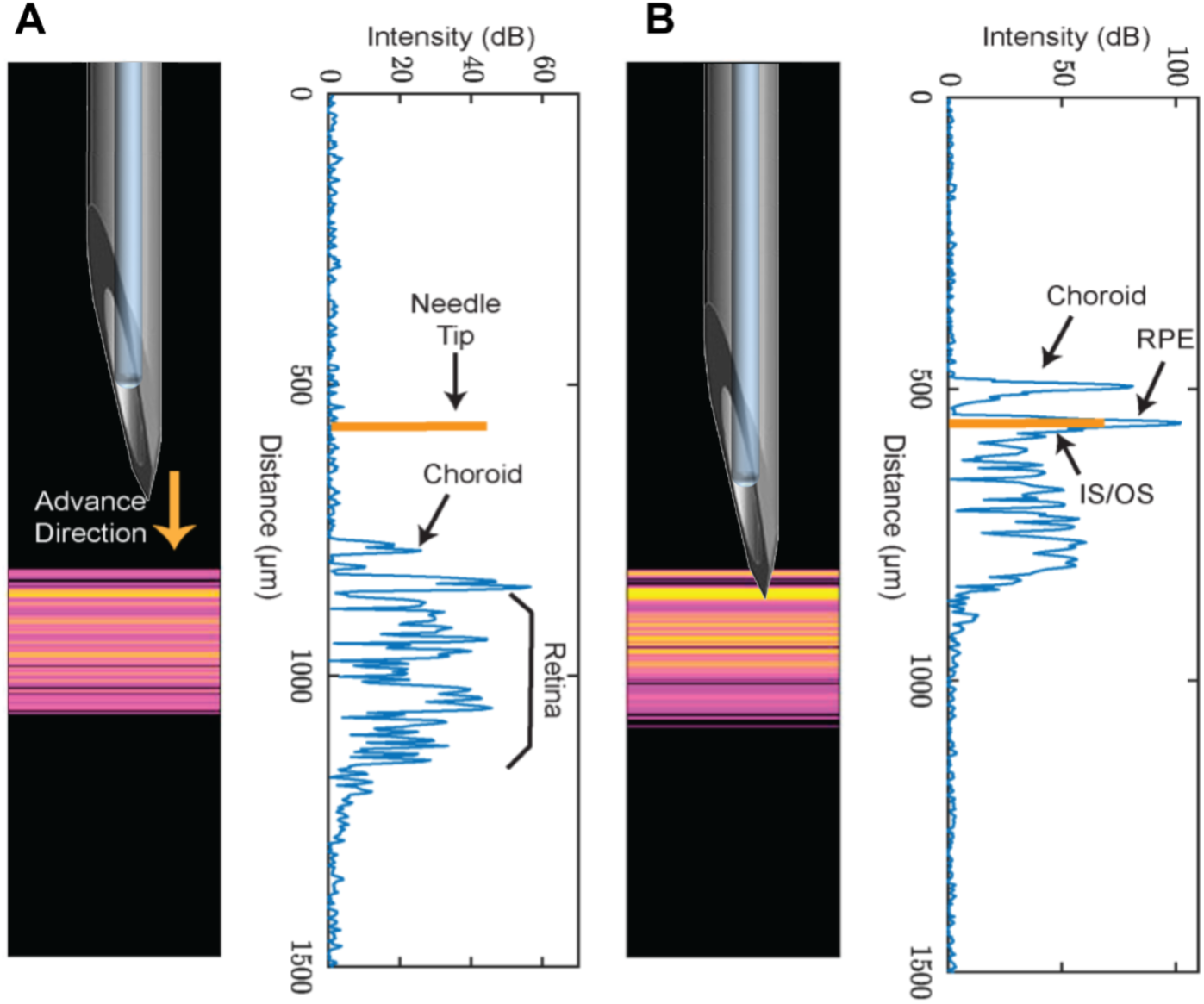
*In vivo* validation of common path swept source optical coherence tomography (CPSS-OCT) based detection of the subretinal plane *ab externo*. A-scan images and corresponding AB-mode images of porcine retina access externally (from the scleral side) as needle advances axially. (A) At ∼200 microns from choroid, (B) needle tip is in the subretinal space. RPE: retinal pigment epithelium. IS/OS: inner

### In vivo implementation of the MISA procedure

The MISA procedure using the CP-SSOCT device, coupled with the COG and SAC components, was then tested on porcine eyes (*n*=4) *in vivo*. We categorized the surgical steps into Phase 1 and Phase 2.

In Phase 1 (**Figure 6**), a rectus muscle is disinserted (a step that is not likely to be necessary in humans) and then the COG was secured on the scleral surface as its vacuum seal was successfully actuated. The 30-gauge needle was successfully advanced through the COG to penetrate the sclera. During this step, OCT imaging showed the depth of penetration in real time. A- and M-mode images were acquired as the CP-SSOCT sensor integrated needle was advanced in the COG through the sclera and into the subretinal space. Hyper-reflective signals presumed to correspond with the laminar depths of the choroid, RPE, and IS/OS layer were identified on the A- and M-mode recordings. The subretinal space was identified as the relatively hyporeflective lamina between the RPE and IS/OS laminae. OCT enabled the identification of the moment then the needle tip had accessed the subretinal space. A subretinal bleb was successfully created by injecting viscoelastic material, as verified by direct visualization. The CP-SSOCT subretinal injection device had enabled the creation a subretinal bleb to gain access to the subretinal space through a transscleral approach, without requiring vitrectomy and trans-vitreal access.

**Figure 6.**
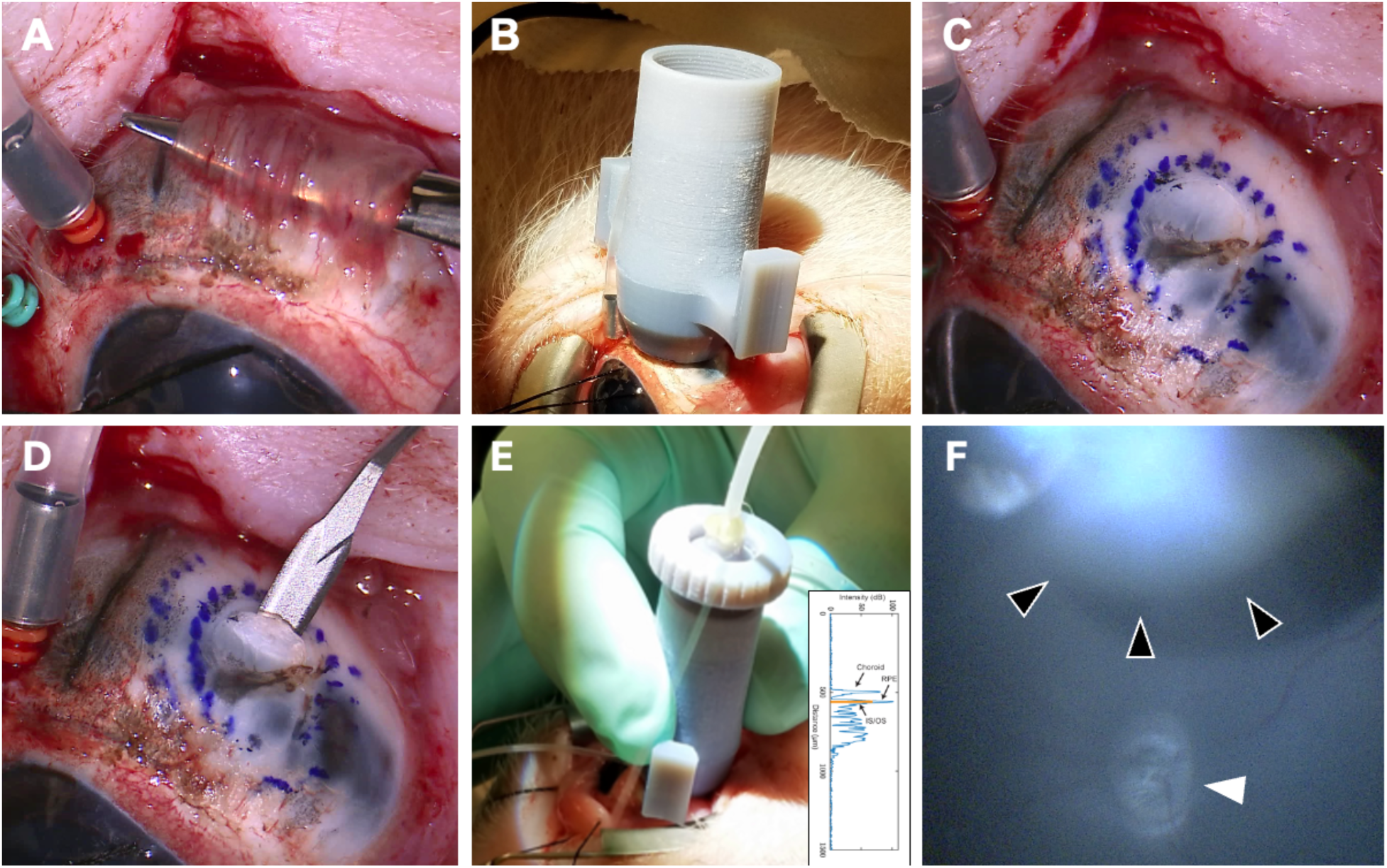
Creating the subretinal bleb during Phase 1 of the minimally invasive subretinal access (MISA) procedure. (A) In the swine model, a rectus muscle is disinserted; however, this step is not necessary in humans. (B) Placement of the coaxial access guide (COG) on the scleral surface. (C) Marking of the inner and outer rings of the COG. (D) Partial thickness sclerotomy (optional step). (E) The COG is secured and a 30-gauge needle is advanced axially towards to eye to penetrate the sclera. During this step, OCT imaging is performed to determine the depth of penetration (inset, same data from Figure 5 shown for illustration purposes). Once the subretinal space is identified by OCT, a subretinal bleb is created by injecting viscoelastic material. (F) The subretinal bleb (black arrowheads) is verified by intraocular visualization (white arrowhead shows the optic nerve head).

In Phase 2, a 5.5 mm sclerochoroidotomy (after diathermy) was created. The SAC was then passed through the sclerochoroidotomy (**Figure 7**) and then axially propagated manually towards the posterior pole. As the SAC was flexible, it easily passed along the subretinal space. When the desired position of the SAC tip in the subretinal space at the posterior pole was achieved, the SAC was sutured to the sclera. The SAC tip was successfully positioned in the subretinal space of the posterior pole in n=4 eyes as confirmed by direct visualization. In the last eye, intraoperative transpupillary OCT (Leica Proveo 8 with EnFocus OCT) imaging of the posterior pole was performed to verify subretinal placement of the SAC tip. The SAC tip was successfully positioned in the subretinal space without injury to the overlying retina in this area (**Figure 7**).

**Figure 7.**
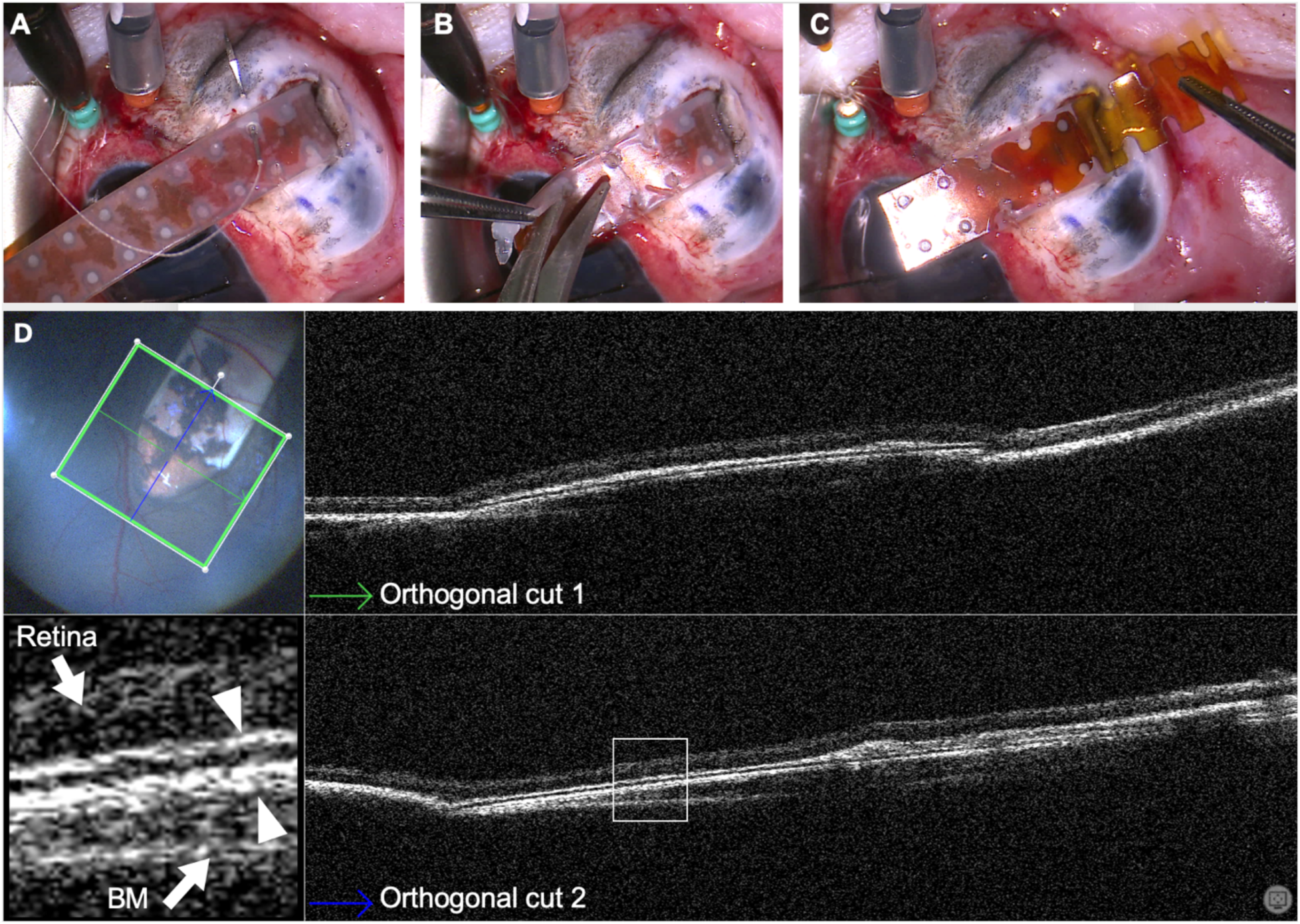
Placing the subretinal access cannula (SAC) during Phase 2 of the minimally invasive subretinal access (MISA) procedure. (A) The SAC tip is advanced through the sclerotomy and secured using a suture once the SAC tip has reached the macula. (B) The latex tube is cut. (C) The top and bottom layers of the SAC are separated to display the SAC lumen. (D) Intraoperative OCT imaging (two orthogonal cuts shown) confirms the placement of the SAC in the subretinal space without retinal injury. The magnified inset shows retina, top and bottom layers of the SAC (arrowheads) and Bruch membrane (BM).

### Complications and mitigating strategies implemented

Complications (**Figure 8**) were classified as major or minor depending on the potential risk of causing severe vision loss. The major complication was retinal incarceration (n=2) and retinal perforation (n=3) at or near the ocular entry site.

**Figure 8.**
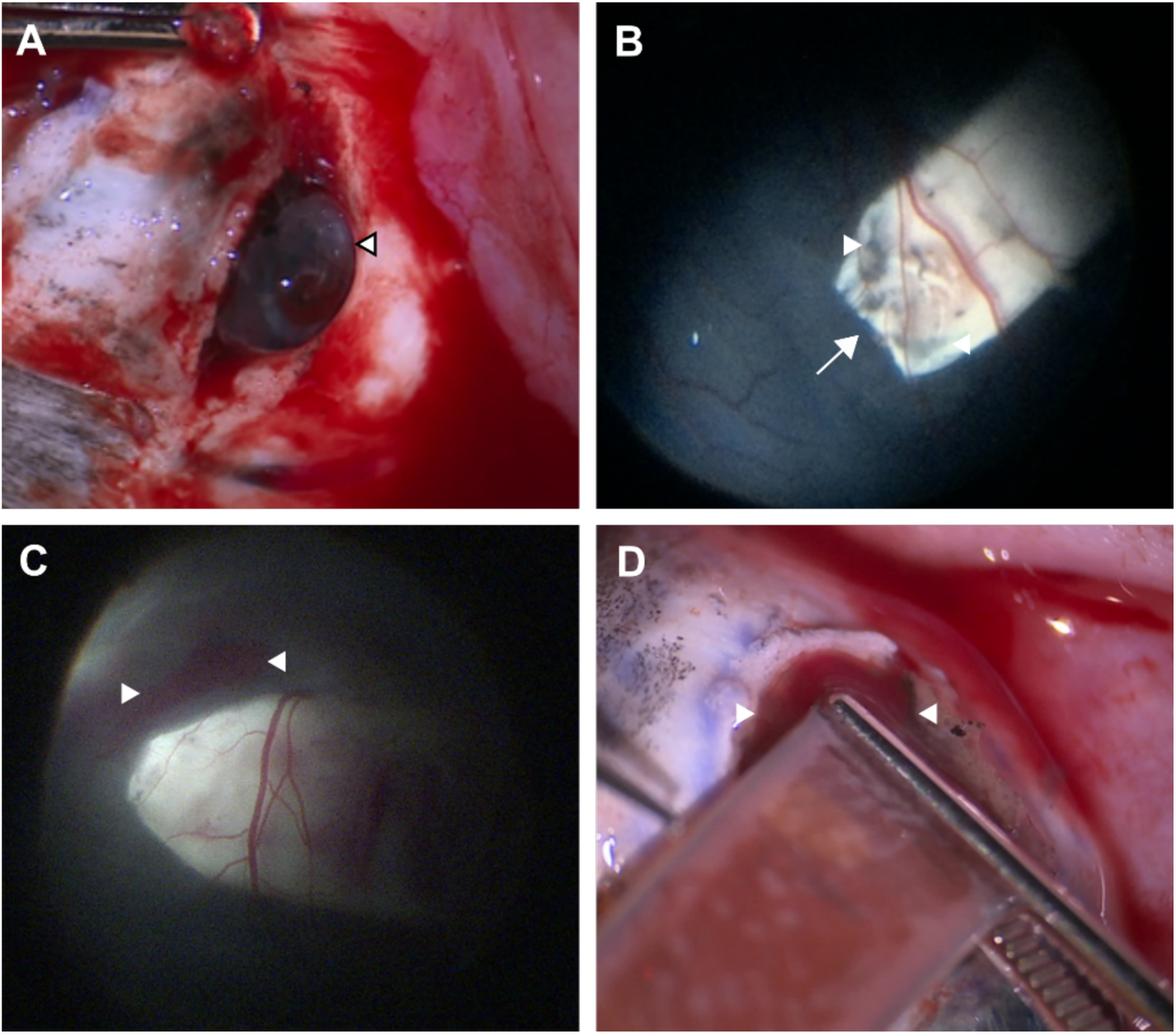
Complications associated with minimally invasive subretinal access (MISA) approach in the swine model. (A) Retinal incarceration through the sclerotomy occurred in the first animal and this was avoided in subsequent cases by turning off the infusion prior to choroidotomy. (B) RPE displacement was seen in one case, where patches of RPE were seen superficial to the SAC tip. (C) Subretinal hemorrhage (arrowheads) was observed in one case that also (D) had choroidotomy site bleeding. The latter complication was avoided in subsequent cases by placing adequate choroidal laser coagulation prior to choroidotomy.

Retinal incarceration in the sclerotomy occurred in the first animal, when intraocular infusion was maintained with the intraocular pressure set 25mmHg during the choroidal incision step. Thus, when the choroid was incised, the injected viscoelastic was expelled and the retina externalized, incarcerated, and ruptured through the sclerotomy/choroidotomy. To mitigate against this risk, we modified the procedure in the next animal such that the infusion flow was reduced (intraocular pressure set at 5 mmHg), however the retina still appeared to bulge outwards upon choroidal incision (without rupture, **Figure 7A**), thus impeding the insertion of the SAC. For all subsequent animals, the infusion was turned off and prior to the choroidotomy step, and we found that the retina no longer bulged out of the eye, there was no retinal incarceration nor perforation, and the viscoelastic was not expelled.

Minor complications were RPE displacement (n=4), mild vitreous hemorrhage (n=3), mild subretinal hemorrhage (n=1) and moderate sclerochoroidotomy site hemorrhage (n=1). No intraocular suprachoroidal hemorrhages were observed.

Vitreous hemorrhage occurred in the first 3 cases presumably because of mechanical stretching of retinal blood vessels by the SAC tip as the initial direction of SAC propagation was not parallel to the plane of the subretinal space. Another possibility was that the hemorrhage occurred because the tip of the chandelier endoillumination or infusion line had touched the retina during ocular manipulation. No retinal breaks were detected along the SAC propagation route. We modified the procedure to ensure that the vector of propagation was angled parallel to the subretinal plane as far as possible, and this adjustment enabled propagation of the SAC to the posterior pole of the pig eye without vitreous hemorrhage in the last animal.

RPE displacement, a minor complication, was observed in one case wherein patches of pigmented RPE cells were observed to lie superficial to the SAC tip as shown in **Figure 8B**.

## Discussion

The data demonstrate proof of concept and feasibility of the MISA procedure. We found that it was possible to place a relatively large planar SAC in the posterior pole of the porcine eye using the CPSSOCT-enabled MISA procedure. To our knowledge, this is the first description of an IGS approach to access a cavity or natural cleavage plane in any body part using fiber-optic distal sensor-based OCT imaging. In the ocular context, the MISA procedure obviated the need for vitrectomy and trans-vitreal access.

Commercially available transpupillary intraoperative OCT allows microscope-integrated OCT head-up visualization system has allowed intraoperative use of OCT to enhance visualization intraoperatively during vitreo-retina surgery[28]. However, uniquely the CPSS-OCT system uses A and M mode images to identify the sub-retinal space from an external transscleral (ab externo) approach.

Initially pioneered for neurosurgery, and now deployed extensively in rhinology and orthopedic surgery, IGS has become the standard of care. Substantial published data indicate that IGS has improved the safety and reduced the invasiveness of numerous surgical procedures in the cranium, nasal sinuses, abdomen, spine, and heart, among others[29]. The subretinal space may join this list of anatomical targets for which morbidity of access could be minimized using an IGS-based approach.

Several other approaches to gain access to the sub-retinal space for cell/tissue-based therapies have been described (recently reviewed in[30]). The first is to deliver cell suspensions into the subretinal space through subretinal cannula via the conventional *ab interno* vitrectomy-based approach[16]. Using this technique, a 38-gauge retinotomy is performed which is presumed to self-seal postoperatively. However, this approach is limited to cell suspension-based therapies, as preformed sheets of cells cannot be passed through a small-gauge lumen without risking cell and scaffold damage. Concern also exists regarding the viability of cells in suspension after being passed through small-gauge devices due to the potential for cell membrane shearing and cell lysis[31]. For the delivery of preformed constructs consisting of cells-and-scaffold[8, 9], a sufficiently large retinotomy is required. Although data on the impact of retinotomy size on surgical adverse events have not been reported, large retinotomies may increase the risk of retinal detachment, epiretinal membrane (ERM) formation and proliferative vitreoretinopathy (PVR).

Another approach involves subretinal delivery at or near the posterior pole through an *ab externo* approach via a long path along the suprachoroidal space[7]. In order to deliver the cell suspension into the subretinal space at the posterior pole, the sclera was exposed and the iTrack model 275 cannula (iScience) was threaded posteriorly through the suprachoroidal space. However, adverse events related to the surgery were reported including retinal perforations and detachments[7]. Recognizing the need for intraoperative imaging at the initial penetration site to avoid retinal penetration, the authors reported that they adapted their technique by introducing an intraocular endoscope into the vitreous cavity after vitrectomy[7]. The MISA approach recognizes the importance of IGS at the time of ocular penetration and bleb creation but leverages *ab externo* distal sensor OCT imaging instead of *ab interno* endoscopy. We believe that this is a simpler approach than endoscopy.

The subretinal space was the anatomical target of the alpha-IMS retinal implant, developed by Eberhart Zrenner and colleagues. The *ab interno* vitrectomy-based approach was used to place the alpha-IMS retinal implant[32]. The alpha-IMS retinal implant was 3.2 mm wide, smaller than the SAC (5 mm width) that we tested. In the alpha-IMS retinal implant cases, pars plana vitrectomy was performed and a peripheral subretinal bleb was created under direct intraocular visualization[23].

Suprachoroidal delivery is distinct from subretinal delivery. The suprachoroidal space can be accessed safely as a non-IGS procedure[18–22]. Published suprachoroidal devices include Oxulumis (Oxular Limited, Oxford, UK) and the SCS Microinjector (Clearside Biomedical[18, 19, 22], Georgia, USA).

We found several major and minor complications that currently limit the clinical implementation of the MISA approach without further optimization. Priorities for future research include improving the graphical user interface of the CP-SSOCT system and adjusting the COG parameters to optimizing the angle of subretinal approach, among others.

## Conclusions

The principle of MISA is to use real-time distal sensing and imaging to enable safe and precise navigation towards, into, and along the subretinal space. System components of MISA include CP-SSOCT, COG, and SAC. Here, in a set of pilot *ex vivo* and *in vivo* experiments at modest scale using component prototypes, we demonstrated proof-of-concept and feasibility of the MISA procedure. We observed major and minor complications that indicate the need for further optimization before MISA can be deployed clinically. With further technical and procedural refinements, the MISA system could be implemented to deliver subretinal therapeutics without the need for vitrectomy or retinotomy, thus reducing the morbidity of surgical access. The MISA procedure appears to be a feasible approach to improve clinical outcomes of cell and gene therapeutics in the eye.

## Ethics

The Johns Hopkins Animal Care and Use Committee approved the research protocol (SW16M257).

## Author contributions

MSS, KW and JUK conceptualized the study. MSS, MM, SHB, DA, KL, SW, SL, KP, KW, JUK were responsible for experiments and data collection. CBT, KW, JUK, KL, SG, and MSS were responsible for data analysis and manuscript writing.

